## Supplemental Information for "Heat Stable and Intrinsically Sterile Liquid Protein Formulations"

Content	Page Number

Materials	S1

Supplementary Fig. 1. Representative C_P_ plots from DSC measurements in PBS	S2

Supplementary Fig. 2. ^1^H-NMR spectra of PFNA:BSA complex	S3

Supplementary Fig. 3. ^19^F-NMR spectra of PFNA:BSA complex	S4

Supplementary Fig. 4. FTIR Amide protein spectra after PFNA interaction	S5

Supplementary Fig. 5. FTIR C–F spectra after PFNA-protein interaction	S6

Supplementary Fig. 6. PFOc-BSA extended bacterial contamination	S7

Supplementary Fig. 7. Full size figure of serologic toxicology results	S8

Table S1: Tabulated serological toxicity results	S9

Supplementary Fig. 8. H&E tissue sections (40X magnification)	S10

**Materials**

n-Perfluorooctane (PFOc) solvent was purchased from Oakwood Chemical (Estill, NC). Perfluorononanoic acid (PFNA) was purchased from Alfa Aesar (Haverhill, MA). 1X Phosphate Buffer Saline (PBS) was purchased from Corning (Corning, NY). Coomassie Protein Assay Reagent 1X, Bovine Serum Albumin (BSA) and o-Nitrophenyl-β-D-galactopyranoside (ONPG) were purchased from was purchased from Fisher Scientific (Hampton, NH). Human Hemoglobin Lyophilized powder, β-Galactosidase from E. *coli* Grade VI (β-Gal), Human Serum IgG, Trypsin from Bovine Pancreas Type I, Proteinase K from T. *album*, 4-methylumbelliferyl β-D-galactopyranoside (µ-Gal), Nα-Benzoyl-DL-arginine 4-nitroanilide hydrochloride (BAEE) were purchased from Sigma-Aldrich (St. Louise, MO). 50% v/v Hydrochloric acid (HCl) was purchased from Aqua solution (Jasper, GA). 100% Bleach solution was acquired from Clorox™ (Oakland, CA). Green Fluorescent Protein (GFP) was a generous gift from Dr. Joel Schneider of the National Cancer Institute (Bethesda, MD).

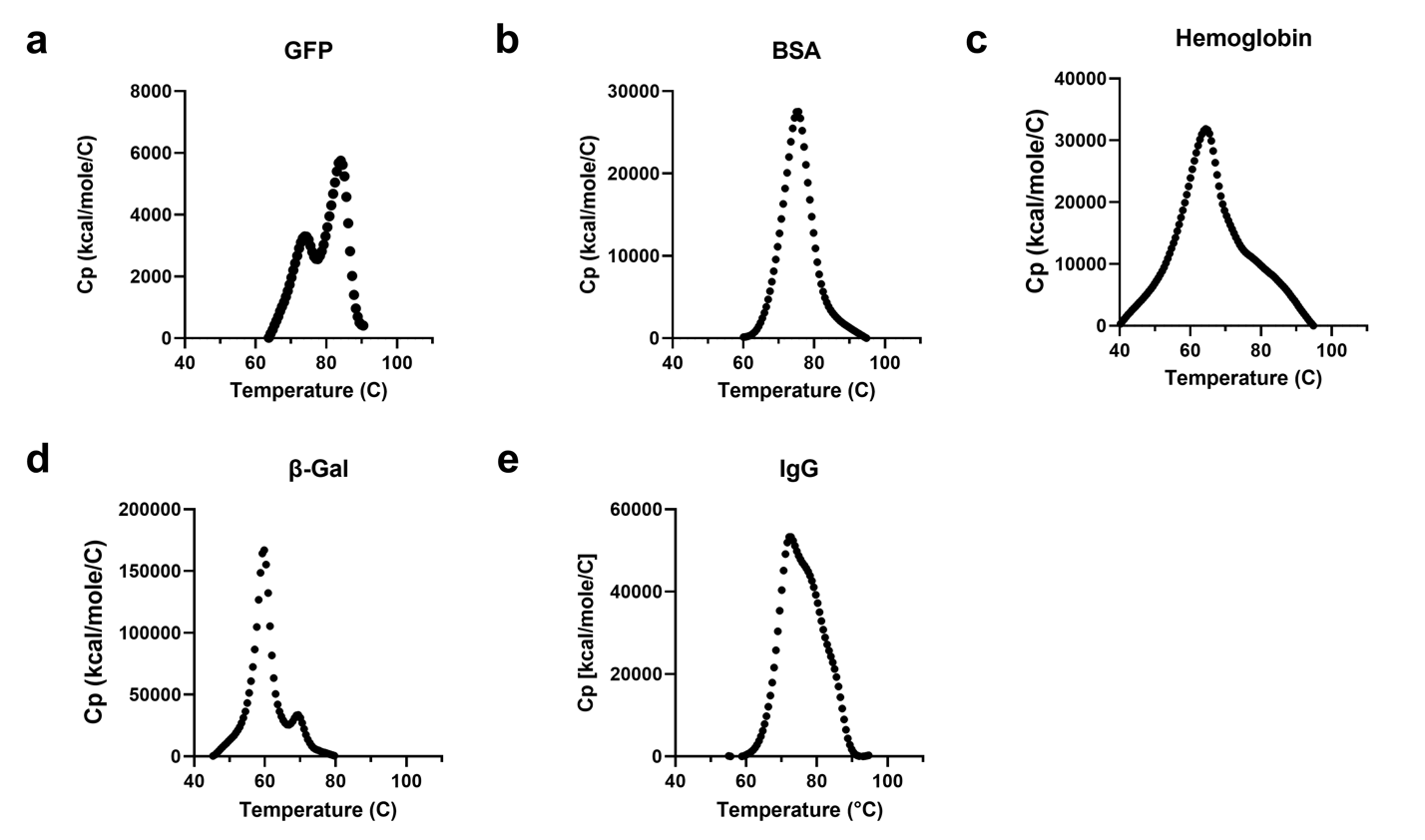

**Supplementary Fig. 1:** Representative DSC heat capacity profiles for (**a**) GFP, (**b**) BSA, (**c**), Hemoglobin, (**d**) β-Gal, and (**e**) IgG in saline.

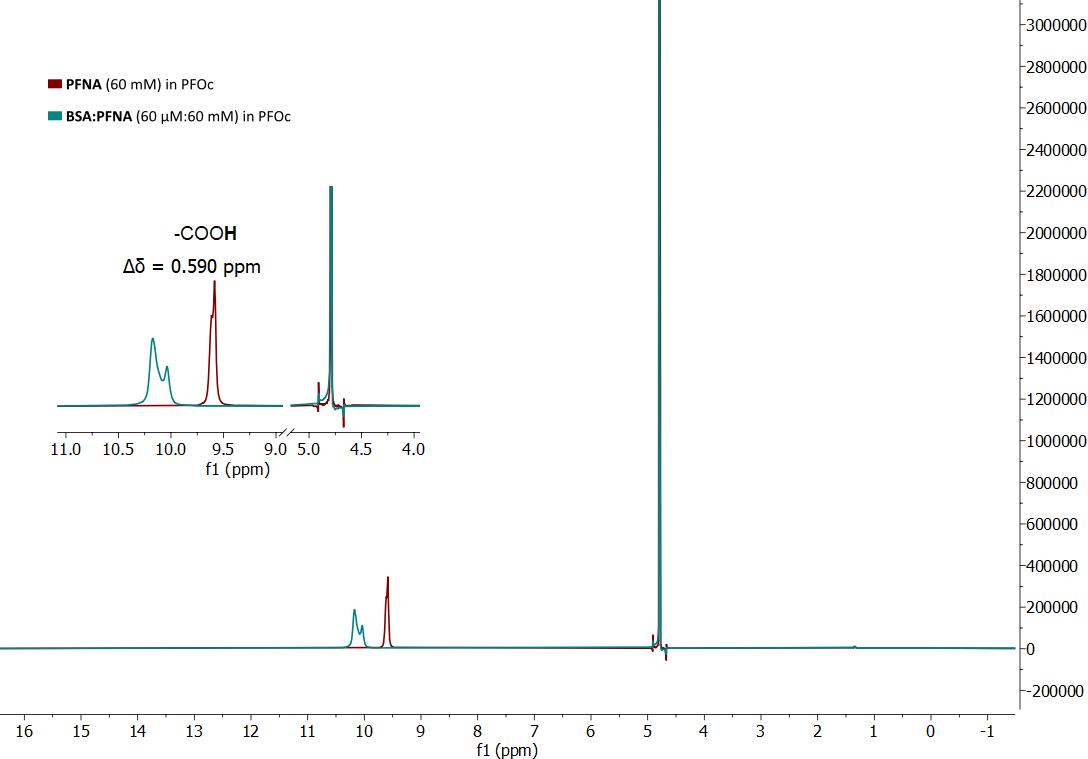

**Supplementary Fig. 2:** ^1^H-NMR spectrum of PFNA (red) or BSA:PFNA complex (blue, 1:1000 molar ratio).

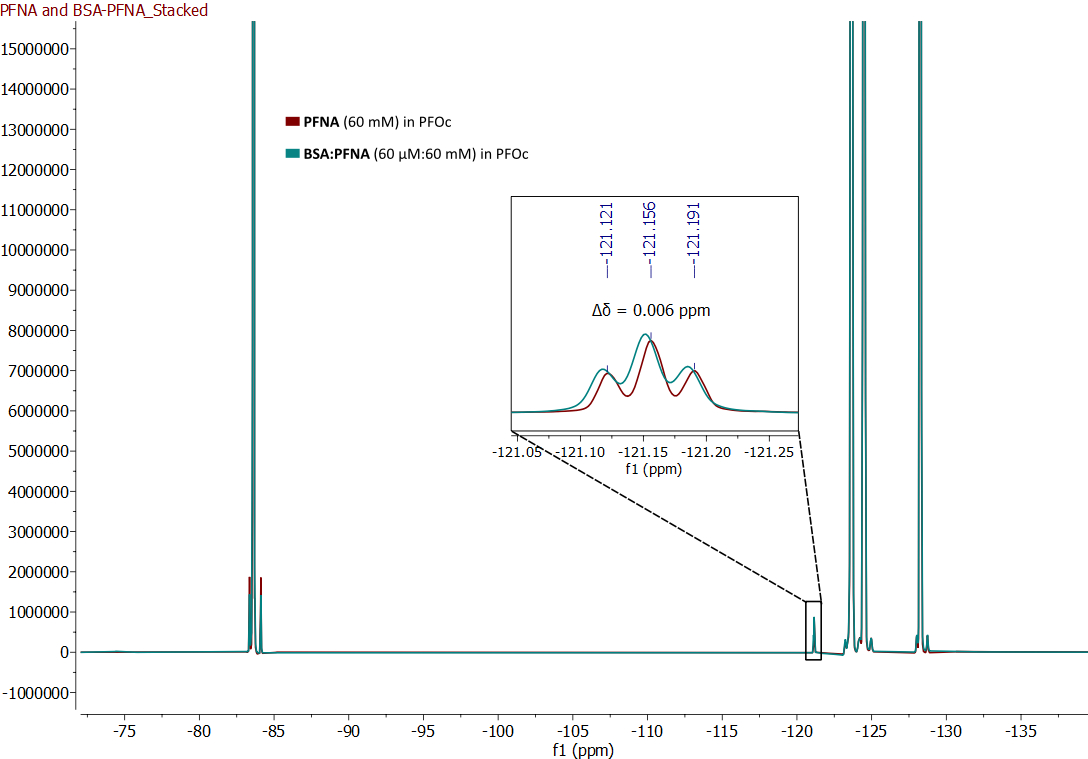

**Supplementary Fig. 3:** ^19^F-NMR spectrum of PFNA (red) or BSA:PFNA complex (blue, 1:1000 molar ratio).

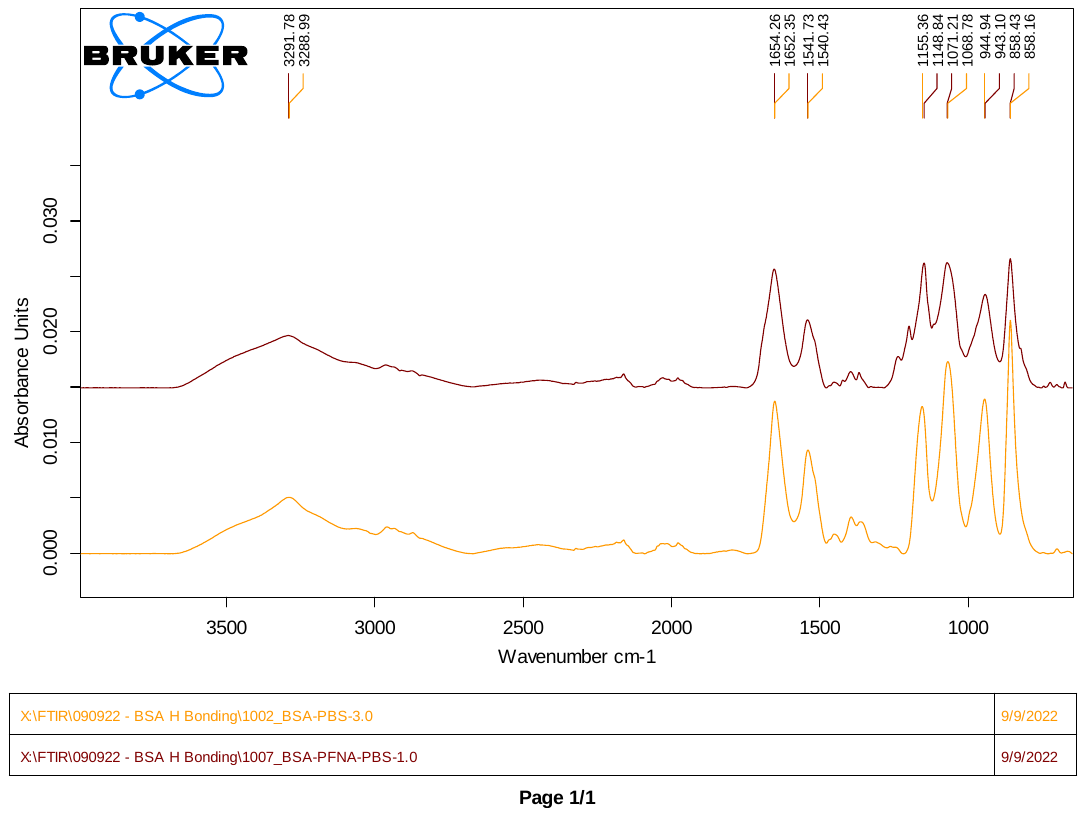

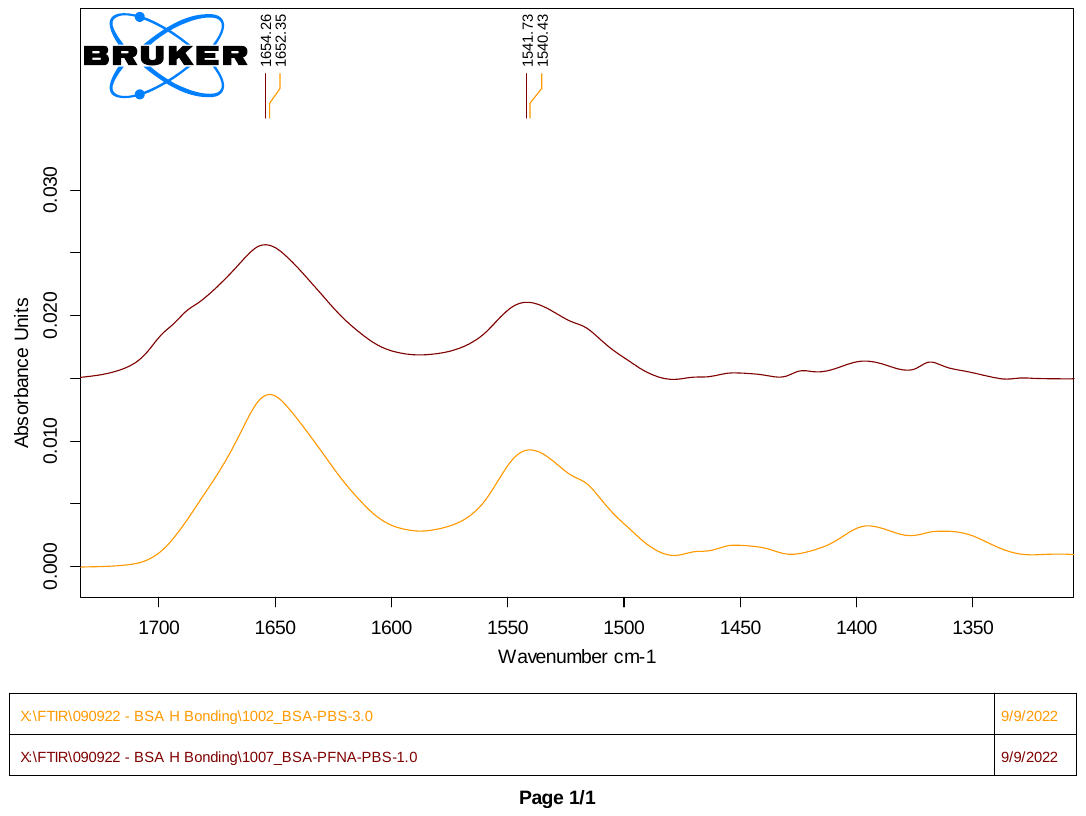

**Supplementary Fig. 4:** FTIR tracing shift in Amide peaks of proteins after PFNA interaction. Spectra are shown in (top) full or (bottom) magnified between 1750 – 1300 cm^-1^. Spectra were taken from lyophilized (orange) free BSA and (brown) PFNA-treated BSA samples.

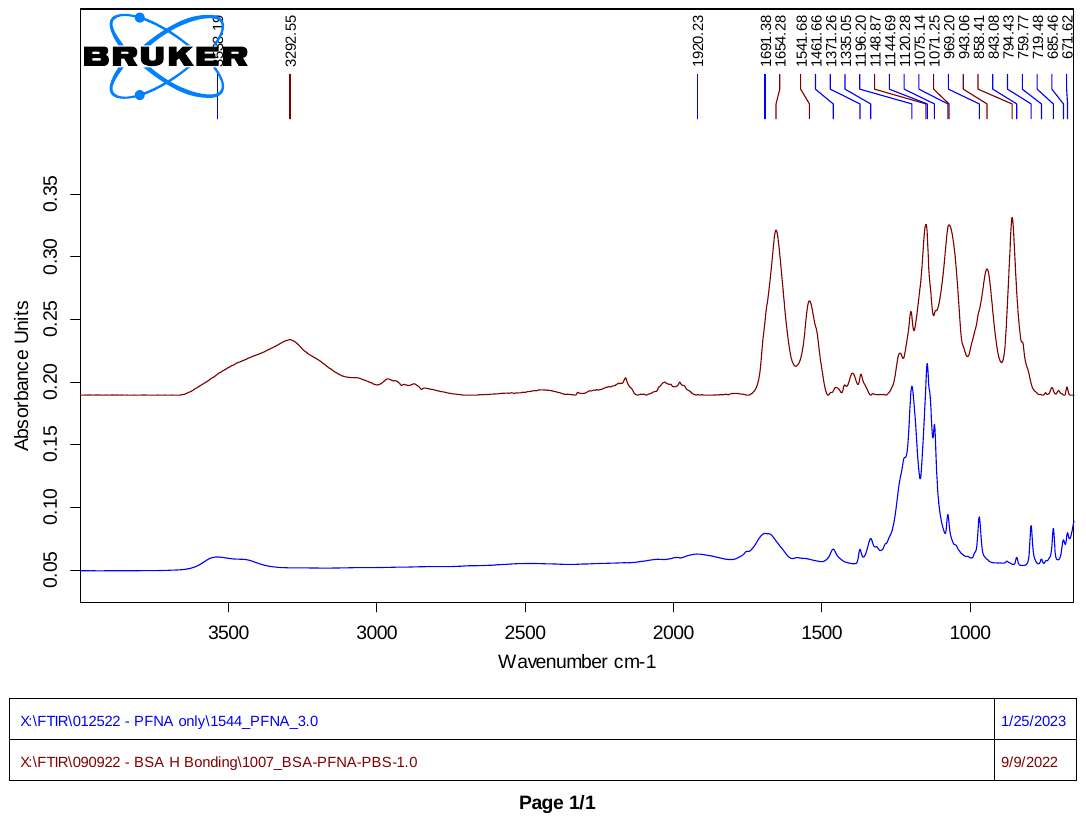

**Supplementary Fig. 5:** Full FTIR spectrum tracing shift in C–F peaks of PFNA after protein interaction. Spectra were acquired from (blue) PFNA or (brown) lyophilized BSA-treated PFNA samples.

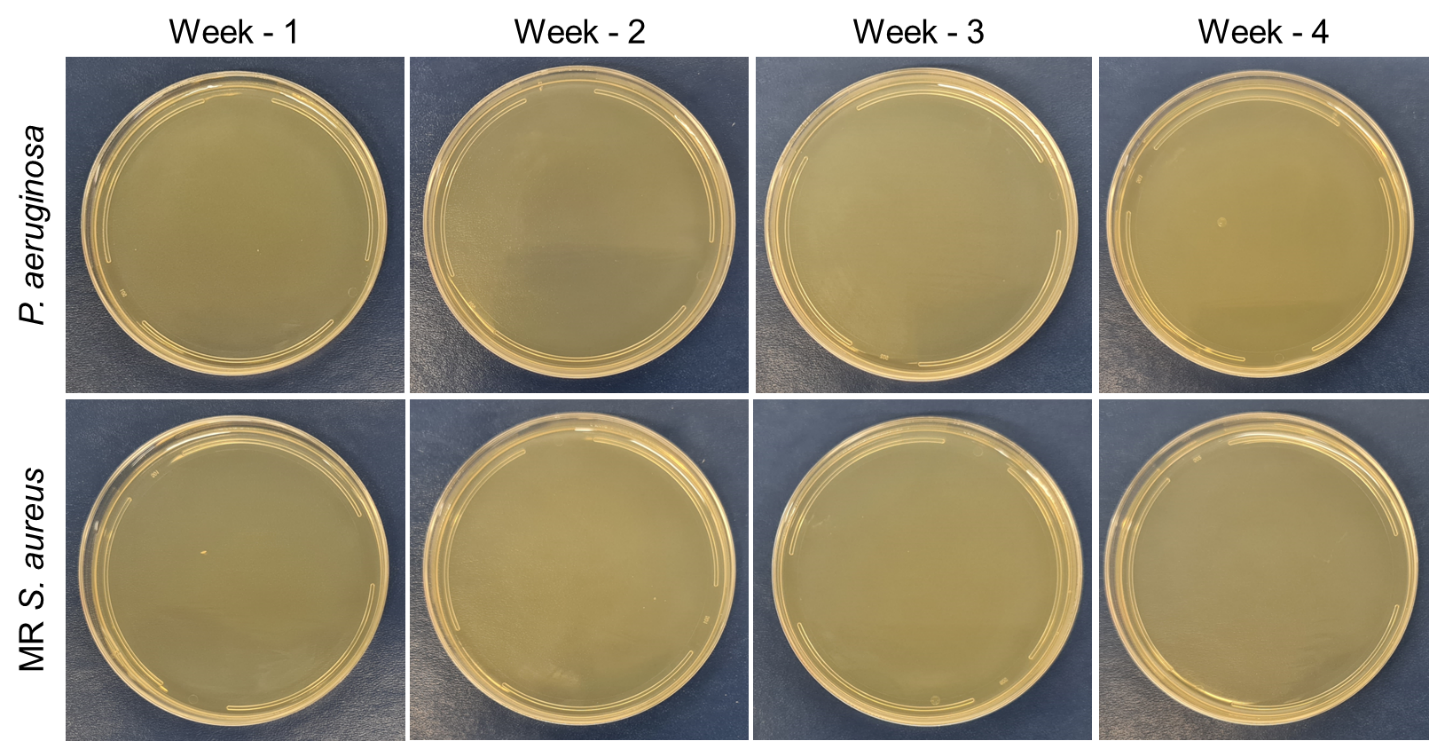
**Supplementary Fig. 6:** Representative optical images of agar plates after addition of PFOc-BSA samples contaminated with (*top row*) *P. aeruginosa* or (*bottom row*) Methicillin resistant *S. aureus* and allowed to culture for 1 – 4 weeks.

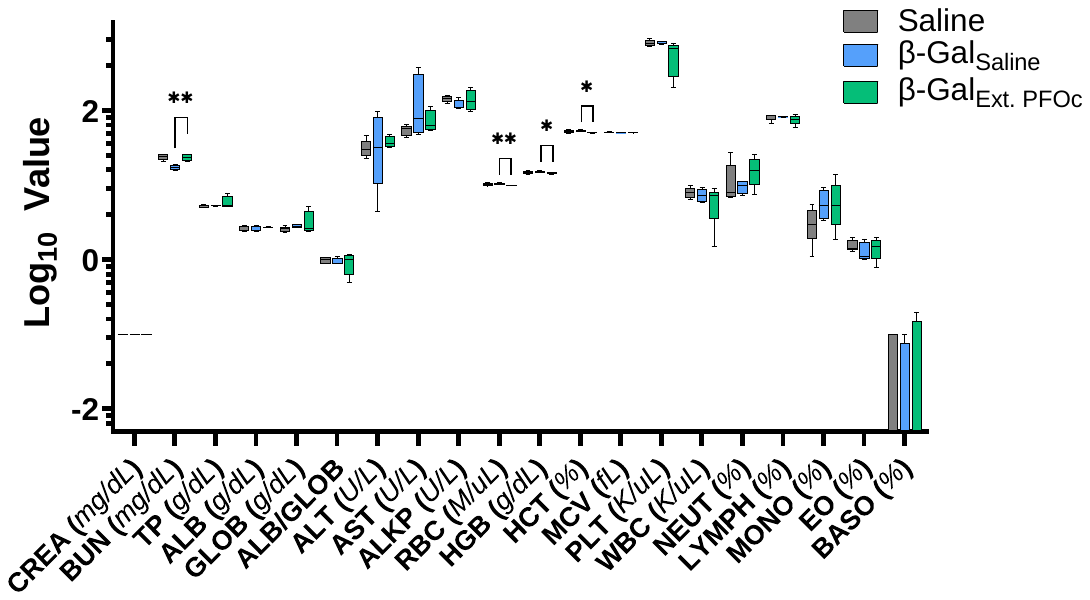

**Supplementary Fig. 7:** Full size figure showing serologic toxicology results from C57BL/6J mice 24 hours after administration of saline (control), β-Gal_Saline_ or β-Gal_Ext. PFOc_. Data shown as box and whisker plot ± s.d. of n = 4-5 technical replicates. Statistical significance determined using Student’s t-test and represented as * p < 0.05; all other comparisons were found not to be significant (p > 0.05).

**Table S1**: Tabulated serological toxicity results

| **Marker** | **Saline** | | **β-Gal_Saline_** | | **β-Gal_Ext. PFOc_** | |
| --- | --- | --- | --- | --- | --- | --- |
|  | *Average* | *Std. Dev.* | *Average* | *Std. Dev.* | *Average* | *Std. Dev.* |
| CREA (*mg/dL*) | 0.10 | 0.00 | 0.10 | 0.00 | 0.10 | 0.00 |
| BUN (*mg/dL*) | 24.00 | 2.12 | 24.00 | 2.65 | 17.50 | 1.29 |
| TP (*g/dL*) | 5.28 | 0.27 | 6.10 | 1.47 | 5.40 | 0.14 |
| ALB (*g/dL*) | 2.62 | 0.22 | 2.73 | 0.06 | 2.58 | 0.22 |
| GLOB (*g/dL*) | 2.58 | 0.22 | 3.40 | 1.56 | 2.85 | 0.13 |
| ALB/GLOB | 1.00 | 0.10 | 0.90 | 0.36 | 0.95 | 0.10 |
| ALT (*U/L*) | 31.40 | 8.73 | 38.00 | 8.72 | 41.63 | 39.83 |
| AST (*U/L*) | 54.60 | 8.20 | 79.67 | 29.02 | 146.75 | 159.15 |
| ALKP (*U/L*) | 145.80 | 13.88 | 139.67 | 57.46 | 120.00 | 19.48 |
| RBC (*M/uL*) | 10.20 | 0.41 | 9.94 | 0.13 | 10.54 | 0.27 |
| HGB (*g/dL*) | 14.90 | 0.56 | 14.53 | 0.25 | 15.15 | 0.47 |
| HCT (*%*) | 52.66 | 2.19 | 50.65 | 0.95 | 53.15 | 1.74 |
| MCV (*fL*) | 51.62 | 0.33 | 50.98 | 0.54 | 50.43 | 0.39 |
| PLT (*K/uL*) | 816.20 | 77.85 | 526.25 | 286.39 | 834.25 | 43.87 |
| WBC (*K/uL*) | 8.06 | 1.32 | 5.44 | 2.76 | 7.42 | 1.60 |
| NEUT (*%*) | 11.98 | 8.67 | 13.73 | 4.74 | 9.65 | 1.99 |
| LYMPH (*%*) | 83.24 | 8.78 | 80.15 | 6.18 | 83.15 | 0.58 |
| MONO (*%*) | 3.20 | 1.63 | 4.45 | 1.98 | 5.90 | 2.70 |
| EO (*%*) | 1.54 | 0.28 | 1.60 | 0.29 | 1.28 | 0.42 |
| BASO (*%*) | 0.04 | 0.05 | 0.08 | 0.10 | 0.03 | 0.05 |

**Supplementary Fig. 8:** Histopathological analysis of organs. 40X magnification of H&E stained lung, kidney, liver and spleen tissue sections from mice treated with saline (control), β-Gal_Saline_ or β-Gal_Ext. PFOc_. Scale bar = 25 µm. Each treatment group consisted of 4 mice and analysis was performed on four random fields at 40X for each organ and analyzed in a blinded manner.

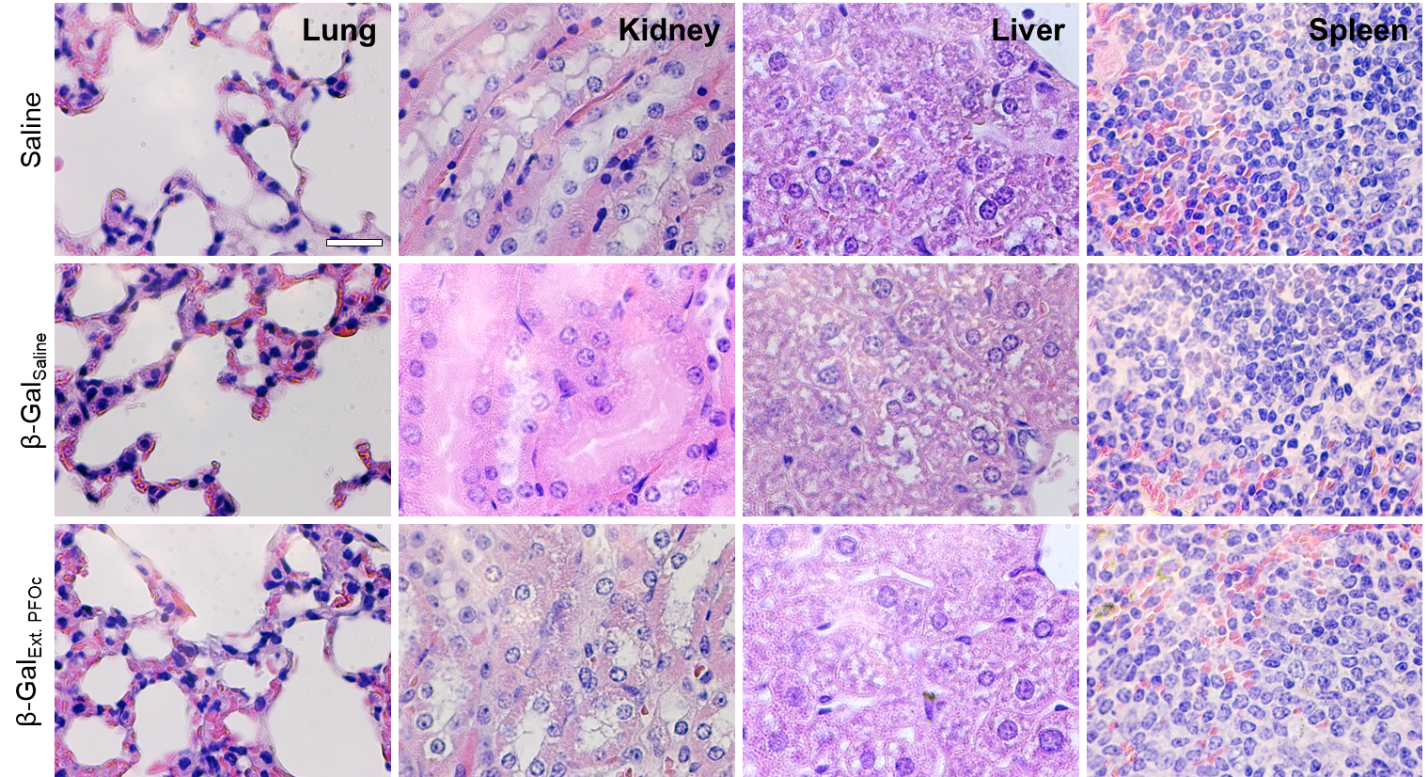
